# Validation and pre-analytical considerations for processing cerebrospinal fluid samples on a high-throughput proximity extension assay platform

**DOI:** 10.1101/2025.04.11.648439

**Authors:** Marieta Hyde, Michael Cunningham, Xiyun Chai, Hengcheng Alvis Hu, Charles Lu, Aparna Vasanthakumar

**Affiliations:** AbbVie Inc., 1 N Waukegan Rd, North Chicago, IL 60064

**Author notes:** Argo Biopharma, 581 Shenkuo Road, 4th Floor, Zhongtong Building, Pudong New Area, Shanghai (Xiyun Chai). Contributed equally (Marieta Hyde, Michael Cunningham). Contributed equally (Charles Lu, Aparna Vasanthakumar). Corresponding authors: Aparna Vasanthakumar, 1 N Waukegan Rd, AP31-1, North Chicago, IL 60064, Charles Lu, 1 N Waukegan Rd, AP31-1, North Chicago, IL 60064.

**Keywords:** Protein profiling, Proximity Extension Assay, Cerebrospinal Fluid, freeze/thaw, Preanalytical considerations, Detectability, Validation, Freeze-thaw cycles

## Abstract

**Background:** Analysis of cerebrospinal fluid (CSF) facilitates the understanding of brain-specific molecular changes that may associate with disease progression. Proximity extension assays (PEA) have been deployed in several CSF studies, however the validation of the assay and impact of freeze-thaw cycles on the protein signal has not been documented. We sought to (1) validate the assay on the PEA platform and (2) evaluate the effect of freeze-thaw cycles on the detectability of analytes on the PEA platform.

**Results:** We have validated the PEA with Next Generation Sequencing (NGS) readout assay and report on the detectability and coefficient of variation observed in CSF samples. We have also evaluated proteomic signals with a minimum of 3 and a maximum of 9 freeze thaw cycles and detected very minimal change in signal with increasing cycle number.

**Conclusion:** Our study is the first to validate PEA using NGS readout platform with CSF samples. We report lower protein detection rates and higher variability in the expansion panels compared to the original 4 panels, with acceptable variation above detectability threshold. In addition, our work demonstrates that the proteomic signal is robust and continues to be stable across multiple freeze thaw cycles. This is highly impactful to the processing and analysis of clinical samples and facilitates the investigation of samples with variable pre-analytical conditions.

## Background

Proteins play a significant role in most biological events in the human body and often are the main subject in understanding disease mechanisms and pathological dysfunctions. Studying the proteome and monitoring changes can help identify novel drug targets, as well as develop diagnostic tests and personalized therapies. In the past, it was a significant challenge to apply proteomic profiling to clinical studies due to several limitations of the technology, such as the number of detectable proteins, sensitivity/limit of detection, precision/reproducibility, accuracy, required sample input, sample complexity, and run costs (1,2). Recent advances in proteomics technologies, including mass spectrometry and high-throughput aptamer- or antibody-based detection methods, have enabled comprehensive biomarker analysis of complex protein mixtures (3–5). A dual antibody-based methods that was commercialized by Olink Proteomics utilizes the Proximity Extension Assay (PEA), developed as a targeted readout of up to 96 biomarkers simultaneously (6,7) which was further developed into PEA with Next Generation Sequencing (NGS) readout platform that measures several thousands of protein biomarkers at once(8).

One of the matrices that is used frequently in neurology studies is cerebrospinal fluid (CSF). CSF is a clear, colorless fluid which surrounds the brain and spinal cord and primarily consists of water, diluted proteins, diverse small molecules, and ions (9). Due to its direct interface with the central nervous system (CNS), CSF has seen extensive use as a source of biomarkers to diagnose CNS diseases and assess the effectiveness of CNS drugs (10–12). High throughput proteomics platforms such as Olink and the aptamer-based technology SOMAScan are increasingly being used to perform expression profiling of large numbers of proteins in the CSF simultaneously (13).

Biofluidic samples collected at biobanks and as part of clinical trials are commonly exposed to multiple freeze-thaw cycles prior to arriving for analysis in proteomics research facilities. Proteins found in CSF as in other matrices are prone to damage and misfolding induced by freeze-thaw cycling prior to assay (13–16). Despite the potential negative consequences, multiple freeze-thaw cycles are frequently unavoidable due to logistical limitations, prolonged experimentation, and re-purposing samples with new technologies or different cohort sets for new biological questions.

Although several studies have applied the NGS based PEA platform to analyze protein profiling with CSF samples (13,16,17), there is currently no validation data available in the literature. We define validation as an evaluation of the assay performance focusing on reproducibility and detectability of the analytes. The only offered information stating the performance of CSF when using PEA is provided by the vendor on their Mouse Exploratory panel, reporting on the percent detectability.(18). Here, we provide comprehensive performance assessments of CSF with PEA assay by using 6 human CSF samples obtained from commercial vendor. We examined the percentage of detectability and reproducibility of all 8 PEA panels assessing the reliability and the limitations of the assay. Further we tested the effect of multiple freeze-thaw cycles on the integrity of the CSF proteome when studied with PEA. While several other preanalytical conditions can result in loss of protein signal (19–21), our study specifically focused on the impact of repeated freeze-thaw cycles, aiming to replicate quick sample aliquoting processes.

## Materials & Methods

### Samples and design

CSF samples collected from six subjects via lumbar puncture were acquired from PrecisionMed, Inc. (Carlsbad, CA). Two donors had a diagnosis of probable amyotrophic lateral sclerosis (ALS), as determined by the revised El Escorial criteria (22) while the other four were controls with no known health conditions (Table 1). PrecisionMed collected 10-12 mL of CSF from each donor into 15 mL conical tubes, centrifuged the tubes at 1200 RCF at room temperature for 10 minutes immediately after collection, then either transferred the supernatant to 50 mL tubes or directly divided it into 1 mL aliquots in 2 mL cryovials. All samples were subsequently transferred to a −80C freezer. Samples transferred to 50 mL tubes were subsequently thawed and divided into 1 mL aliquots in 2 mL cryovials before being re-transferred to a −80C freezer. These samples are labeled in Table 1 as having experienced one or two freeze-thaw cycles at the vendor, respectively.

**Table 1:** Patient cohort used for validation and freeze/thaw (F/T) cycle analysis.

| DONOR ID | DIAGNOSIS | SEX | AGE AT SAMPLING | COLLECTION DATE | F/T CYCLES AT VENDOR | F/T CYCLES USED IN VALIDATION | MAXIMUM TOTAL F/T CYCLES |
| --- | --- | --- | --- | --- | --- | --- | --- |
| 1 | Probable ALS | M | 66 | 8/7/19 | 2 | 3 | 9 |
| 2 | Probable ALS | M | 62 | 9/16/21 | 2 | 3 | 9 |
| 3 | Healthy control | M | 60 | 12/15/20 | 1 | 2 | 8 |
| 4 | Healthy control | M | 63 | 9/26/19 | 1 | 2 | 8 |
| 5 | Healthy control | F | 63 | 1/8/21 | 2 | 3 | 9 |
| 6 | Healthy control | F | 76 | 8/24/20 | 1 | 2 | 8 |

### Validation

Samples used for the validation part of the study had 4 technical replicates that were not subjected to any additional freeze-thaw cycles at our lab other than those inherent in the PEA assay process and thus had a total of 2 or 3 freeze-thaw cycles, respectively (Table 1).

### Freeze/thaw

In order to simulate the normal circumstances of using clinical samples in day-to-day lab operations, each aliquot was then subjected to 6 additional freeze-thaw cycles prior to PEA analysis. Freeze-thaw cycles were induced by removing samples from −80C storage and incubating them at room temperature for 20 minutes, which was visually confirmed to be sufficient to induce full thawing of all samples. The samples were then re-frozen by transfer to a −80C freezer. One cycle was conducted per day to ensure complete re-freezing. As the CSF begins as liquid and is assayed as a liquid, we defined the number of cycles as the number of times the fluid has been frozen in-between. Thus, a sample collected by lumbar puncture, aliquoted and frozen, then thawed for analysis would be considered to have gone through one freeze-thaw cycle.

### Proximity extension Assay sample processing and data analysis

The CSF samples, including ALS patients, controls, technical replicates, and samples with various freeze/thaw cycles (n=88) were randomized, plated in 2 individual 96-well plates, and frozen introducing the final freeze/thaw cycle (2-3 and 8-9 freeze/thaw cycles used in for validation and freeze/thaw cycles analysis respectively). PEA was performed with Explore 3072 assay based on manufacturer’s protocol (Olink Bioscience, Uppsala, Sweden) using the alternative dilution program to prepare the incubation. The four original panels (Cardiometabolic, Inflammation, Neurology and Oncology) were processed together using the first 96 well plate aliquot, and the remaining four expansion panels (Cardiometabolic II, Inflammation II, Neurology II and Oncology II) were treated collectively several days later with the second 96 well plate aliquot. The constructed libraries we sequenced on a NovaSeq 6000 (Illumina, San Diego, USA) using SP kit according to Olink Bioscience sequencing protocol (Olink Bioscience, Uppsala, Sweden). The generated count files were uploaded to Olink NPX Explore Software V1.0 (Olink Bioscience, Uppsala, Sweden) where the Normalize Protein Expression (NPX) values were generated as a final assay read-out. All statistical analysis on the Olink NPX Explore Software output was carried out using a custom R script using standard R library in v4.2 supplied by Posit. Comparison of the expression level of proteins previously observed to be affected by number of freeze/thaw cycle was performed using a paired t-test

## Results

### Validation of CSF on PEA platform

The first aim of our study was to validate the use of the NGS based PEA platform with CSF by identifying the presence of robustly detectable proteins in each of the 8 panels as well as assessing assay variabilities by comparing samples replicates. Protein detectability across a panel was defined as a percentage of protein (assays) whose NPX values are greater than the limit of detection (LOD) as determined by negative controls on the plate. This metric is useful in that it is a good estimate of proteins with quantifiable expression level on the PEA platform. Figure 1 shows that across the original 4 panels, the protein detectability is all above 60% with the Cardiometabolic panel having the highest value (90%) and Oncology having the lowest value (61%). For the 4 expansion panels, the protein detectability was much lower (58% for Inflammation II and 22% for Oncology II). In fact, we observed that overall, the values for the 4 Expansion panels were all much lower than those of the original panels with differences that can reach as high as 50% (Cardiometabolic vs Cardiometabolic II).This reduction has been observed with plasma samples described by the vendor in an Application note (23) but is more pronounced in CSF. We also assessed the reproducibility of all 8 panels by comparing the Coefficient of Variation (CV) across 18 samples (6 subjects each with 3 identical replicates). Table 2 shows the median CV for each panel across the 6 subjects. With the exception of the Inflammation panel, the CVs for all expansion panels were higher than that of the original panel. We hypothesize that the higher variability in the expansion panel could be due to the higher number of proteins whose NPX value is lower than LOD. We then recalculated CV, this time limiting to only proteins whose NPX values > LOD. Supplementary Table 1 shows the new CV values to be less than 15% on average. This value is comparable to the CV calculated from plate controls and is well within the vendor’s recommendation. More importantly, the mean differences in CV between the original and expansion panels are also reduced (Table 2 and Supplementary Table 1).

**Table 2.**
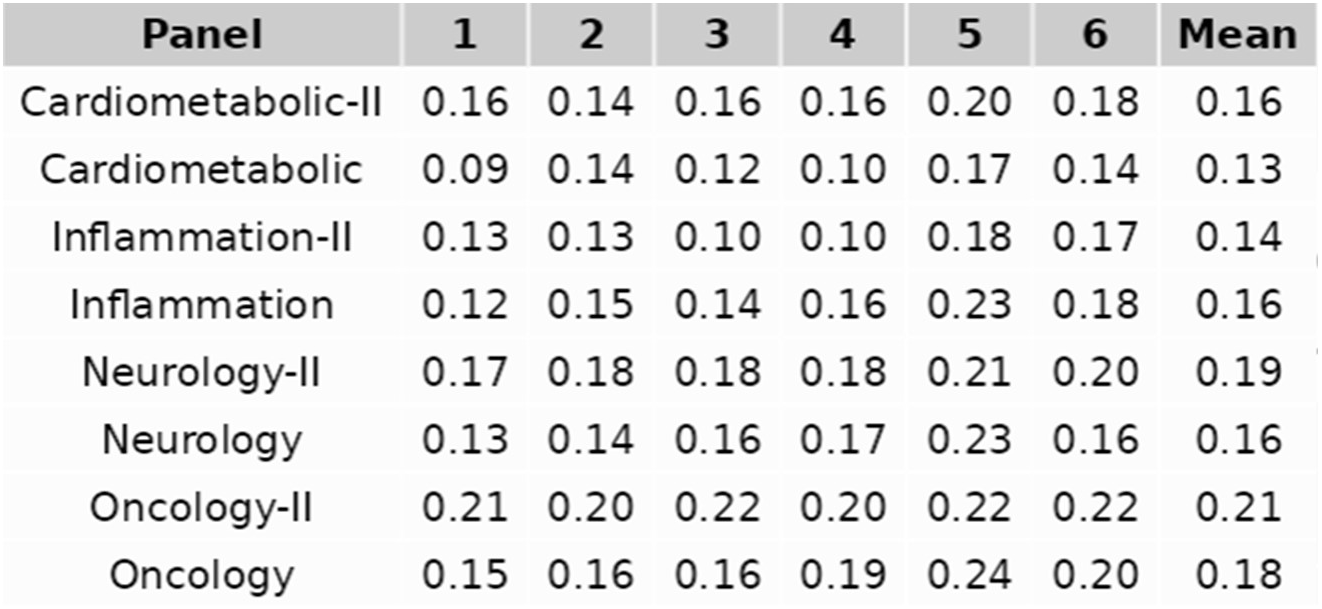
Median Coefficient of Variation per Panel. Includes assays below the Limit of Detection LOD.

| Panel | 1 | 2 | 3 | 4 | 5 | 6 | Mean |
| --- | --- | --- | --- | --- | --- | --- | --- |
| Cardiometabolic-II | 0.16 | 0.14 | 0.16 | 0.16 | 0.20 | 0.18 | 0.16 |
| Cardiometabolic | 0.09 | 0.14 | 0.12 | 0.10 | 0.17 | 0.14 | 0.13 |
| Inflammation-II | 0.13 | 0.13 | 0.10 | 0.10 | 0.18 | 0.17 | 0.14 |
| Inflammation | 0.12 | 0.15 | 0.14 | 0.16 | 0.23 | 0.18 | 0.16 |
| Neurology-II | 0.17 | 0.18 | 0.18 | 0.18 | 0.21 | 0.20 | 0.19 |
| Neurology | 0.13 | 0.14 | 0.16 | 0.17 | 0.23 | 0.16 | 0.16 |
| Oncology-II | 0.21 | 0.20 | 0.22 | 0.20 | 0.22 | 0.22 | 0.21 |
| Oncology | 0.15 | 0.16 | 0.16 | 0.19 | 0.24 | 0.20 | 0.18 |

**Figure 1.**
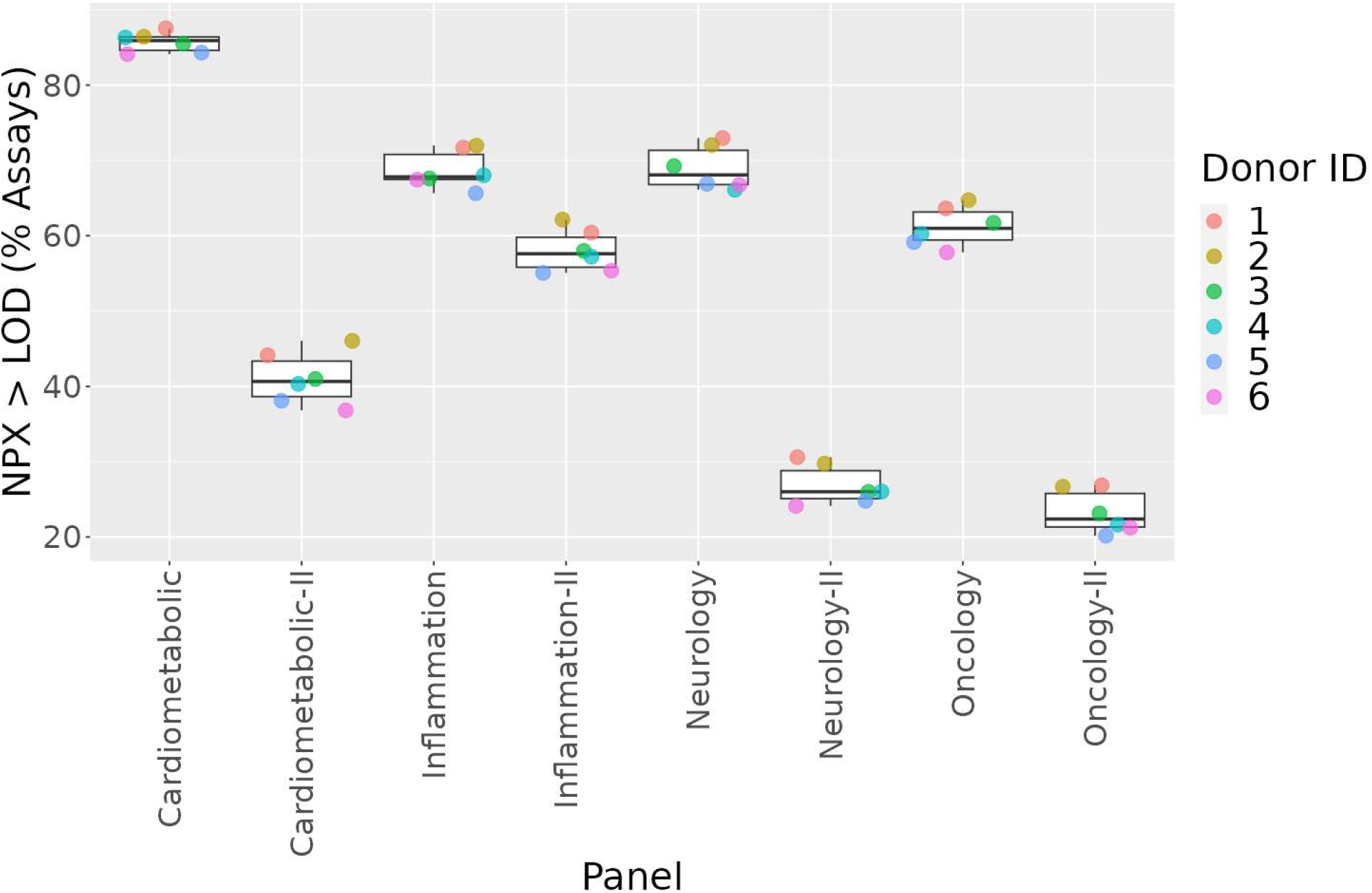
Detectability of Proteins Across Panels. Protein Detectability, defined as % of proteins with NPX values greater than the LOD (limit of detection) shown in box and whisker plots across all 8 panels. Each donor is depicted by colors as indicated in the legend.

### Impact of Freeze/Thaw Cycles on CSF Signal

Having established sample reproducibility and protein detectability for all panels, we next focused on the effect of the number of freeze/thaw cycles on sample quality. For the 6 donors that we have data across 6 different freeze/thaw cycles, we first see if increasing the number of cycles would lead to gross deterioration of the sample and thus reduce the number of detectable proteins relative to baseline. Figure 2 shows that across all 8 panels the number of freeze/thaw cycles does not appear to affect protein detectability. We next assess sample similarities across 36 samples (6 subjects, each with 6 different number of freeze thaw cycles) by PCA plots. Figure 3 shows 8 PCA plots for each of the 8 panels. We see that all the samples primarily cluster by donor instead of number of freeze/thaw cycles (Supplementary Fig. S1). This suggests that the primary difference between all the samples can be explained by the subjects. Although this trend is observed in all 8 panels, unambiguous clustering by subjects is clearly observed in the 4 primary panels. In the 4 expansion panels with higher level of noise, clustering by subjects becomes less pronounced. However, in all 8 panels we do not see any consistent separations between number of freeze thaw cycles across all 6 subjects.

**Figure 2.**
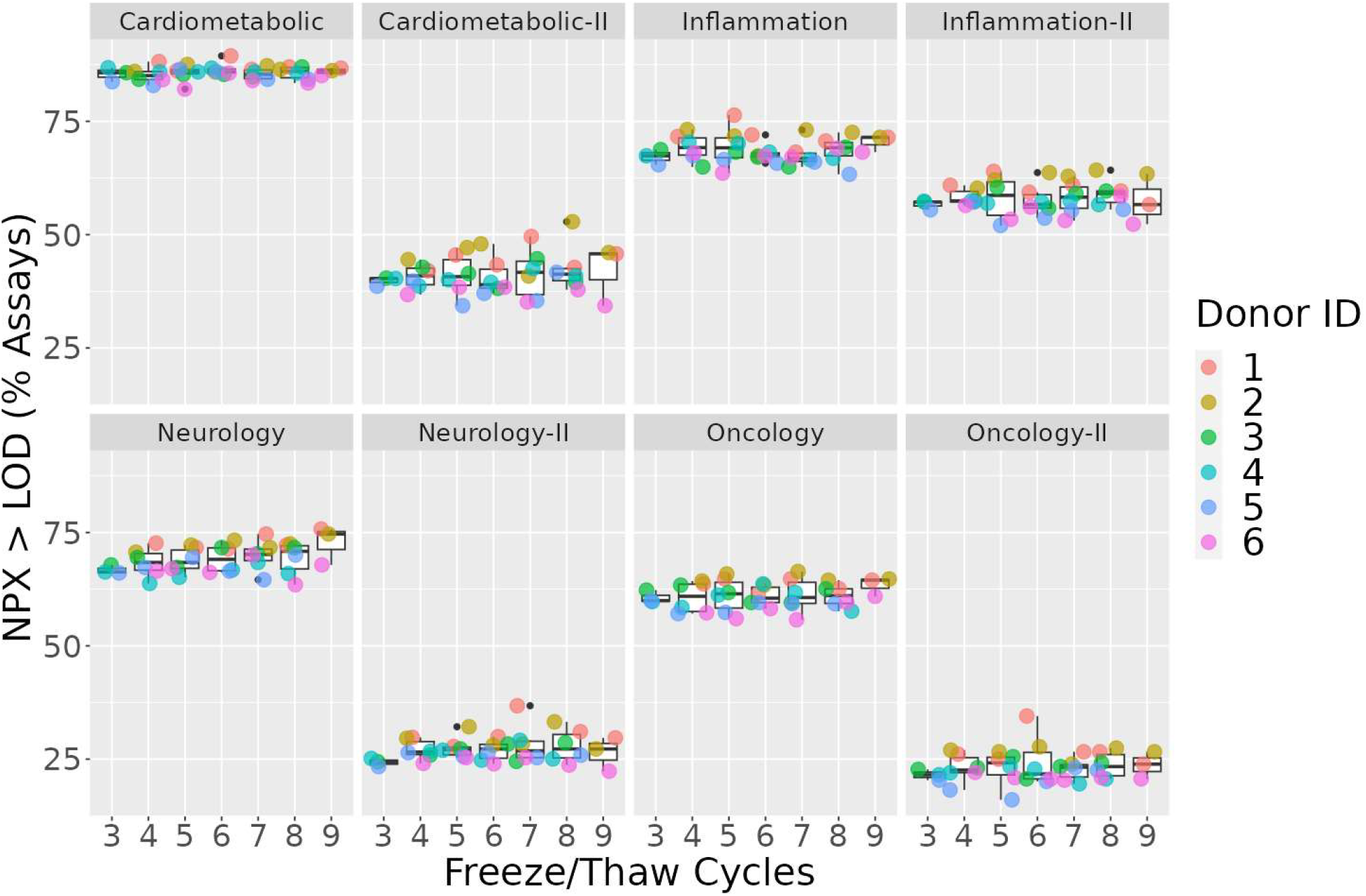
Detectability Vs. Freeze/Thaw Cycles. Protein detectability (NPX >LOD) shown in box and whisker plots across 3-9 freeze/thaw cycles for each individual panel. Subjects are indicated by different colors.

**Figure 3:**
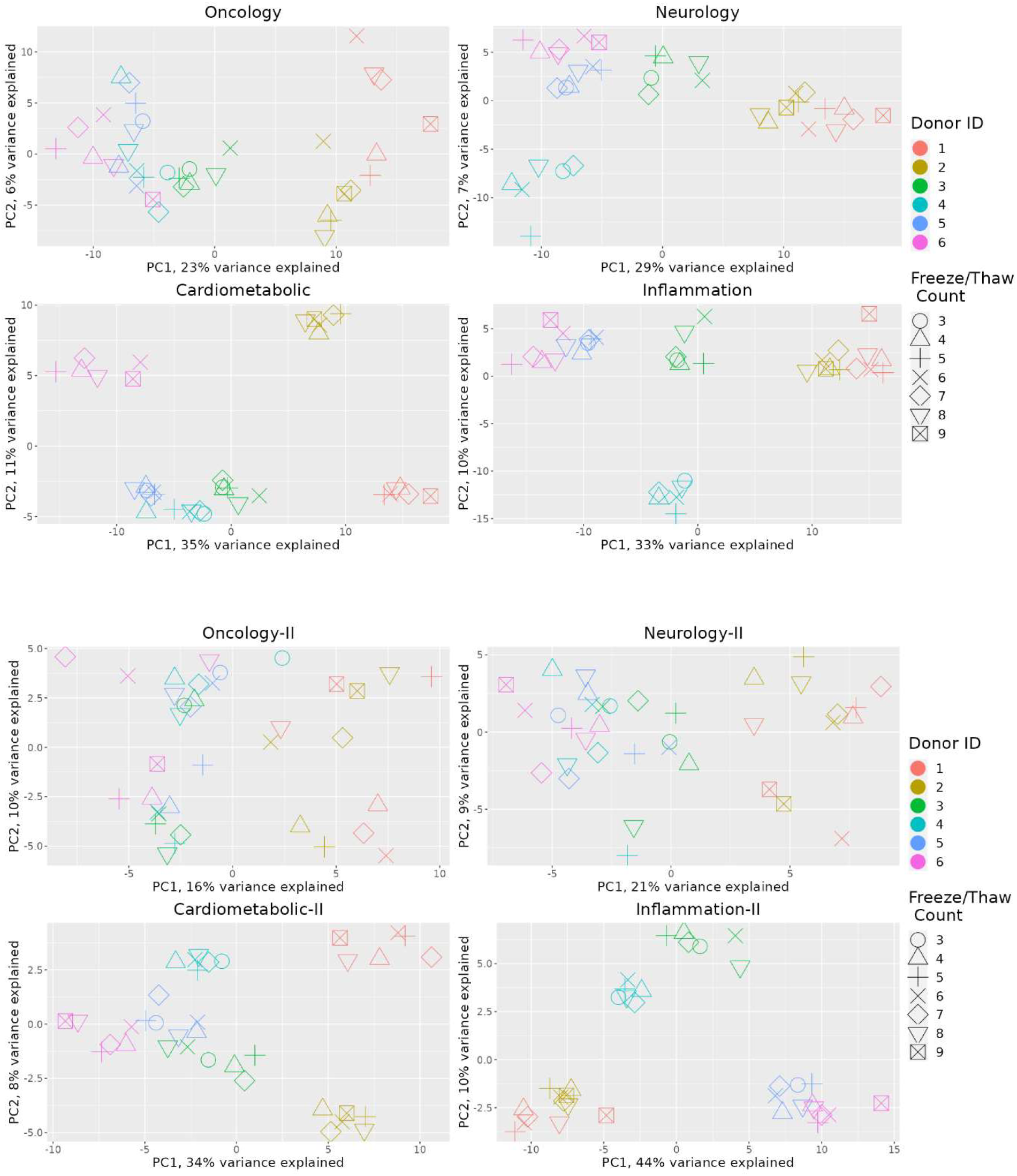
PCA plots by freeze-thaw cycle and stratified by individual panel. Each PCA plot represents an individual panel where donors and freeze/thaw cycles are indicated by colors and shapes respectively.

A previous study (16) using a combination of different proteomic platforms implicated 12 proteins whose expression level was affected by number of freeze thaw cycles when processed with PEA (Supplementary Table 2). We compare the expression level of these proteins in our data for freeze/thaw cycle 4 and 8. Figure 4 shows a paired t-test between cycle 4 and cycle 8 from 6 subjects. None of the 12 proteins show significant differences in expression between the 2 freeze thaw cycles (all of the p values are > 0.05). We then performed a global differential expression test between cycle 4 and cycle 8 with subject as a random effect and we detected no protein with significant differences between the 2 cycles Supplementary Figure. S2.

**Figure 4:**
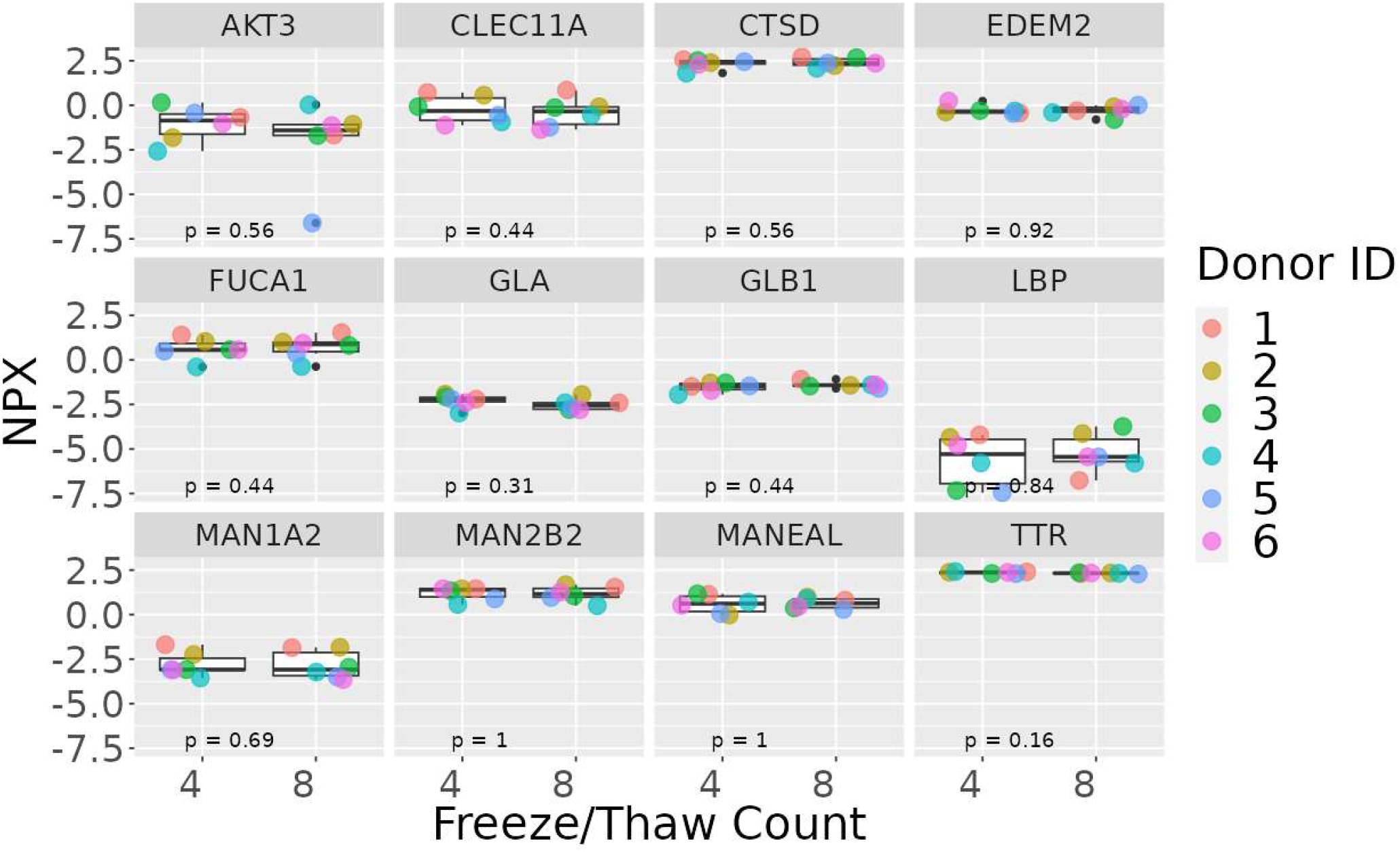
Expression comparison of proteins reported in Hok-A-Hin *et al*. study to be affected by the number of freeze/thaw cycles. P-values for paired t-test are displayed in each facet. NPX values are not found to be significantly different (p < 0.05) between freeze/thaw cycles for any of these assays.

## Discussion

In this study, we examined the performance of PEA in analyzing CSF samples by assessing the percentage of target proteins detected and the reproducibility of the data across replicates. To our knowledge, this is the first study to report on the detectability and reproducibility in CSF using high throughput PEA platform targeting nearly 3000 biomarkers. Our findings indicate that there is a lower percentage of proteins detected and a higher coefficient of variation in the expansion panels (Cardiometabolic II, Inflammation II, Neurology II, and Oncology II) compared to the original four panels (Cardiometabolic, Inflammation, Neurology, and Oncology). These observations can be explained by the nature of the proteins targeted in the different panels where the 4 original panels target extracellular and circulating proteins and the 4 expansion panels cover a higher number of intracellular and disease specific proteins (24). We also found that, when limiting our analysis to proteins with NPX values above LOD, the coefficients of variation fall within acceptable ranges, suggesting that the variability is driven by low-abundance proteins. These results are consistent with previous findings using plasma samples (25,26).

We also investigated the impact of freeze/thaw cycles on protein detectability in CSF. Surprisingly, varying the number of freeze/thaw cycles from 3 to 9 did not have a significant effect on protein detectability across all 8 panels. We demonstrate that the primary variance is driven by donor and not the number of freeze/thaw cycles. Similar findings have been reported previously in a study using the PEA platform that evaluated the stability of proteins in plasma and synovial fluid (27,28). In contrast, another study exploring the impact of freeze-thaw cycles on the CSF proteome that utilized advanced technologies such as slow off-rate modified aptamer and PEA technologies have found that 85% and 76% of proteins respectively remained stable after eight freeze-thaw cycles (16). We closely examined overlapping proteins that were analyzed using PEA between the Hok-A-Hin *et al*. study and ours and we did not observe any significant differences in expression of the proteins between the different freeze/thaw cycles. A few key differences exist between our study and that of Hok-A-Hin *et al*. One distinction is the technique used for conducting the freeze/thaw cycles, where our experiment design focuses on testing conditions for quick sample aliquoting, as Hok-A-Hin et al. explored conditions that involve longer lab processing with extended exposure of the samples at room temperature. Another difference between our study and the Hok-A-Hin *et al*. work is that their group used PEA with qPCR readout and our study used PEA with NGS readout. Even though both approaches use PEA technologies, Olink Proteomics considers them as separate platforms and does not advise them to be used interchangeably. In addition, it is worth considering that there may have been changes in antibody design between the two studies, given the 4 years between ours and the Hok-A-Hin *et al*. study. This discrepancy highlights the necessity for thorough medium and assay-specific determination of the effects of freeze-thaw.

We recognize that sample degradation due to freeze-thaw may occur in first several cycles; however, we believe our results are a robust representation of the changes induced by subsequent cycles. Obtaining unfrozen samples for analysis is not a practical scenario, given that samples from clinical trials or large consortia have typically undergone at least 3 freeze/thaw cycles. In addition, an important detail to highlight is that our study specifically focuses on an older demographic, as the donor samples used were all from individuals aged 60 and above.

Our findings suggest several guidelines for conducting future PEA experiments with CSF. First, differential expression analyses using PEA should be restricted to proteins with NPX values above LOD, as we find that only these likely represent real signal and consistently maintain a CV within the range recommended by vendor. In practice, this means that many proteins, particularly for the expansion panels, will be effectively undetectable in CSF. We also find that detectability does not appreciably decline with additional freeze-thaw cycles, implying that researchers should not regard data collected at up to 8 and potentially more cycles as likely to be substantially compromised. As we cannot rule out that significant loss of signal occurs within the first several cycles, researchers who are able to limit freeze-thaw cycles to fewer than 3 may benefit from doing so; however, it is probable that this will rarely be the case in practice. Finally, as our study conflicts with Hok-A-Hin *et al*. as to which individual proteins remain stable after multiple freeze-thaw cycles, researchers should remain agnostic as to which proteins may undergo degradation with additional cycles, and not consider any particular proteins as more likely to be safe or compromised. Since this work was conducted, the company has released a new, broader kit utilizing PEA technology with NGS readout, targeting more than 5,000 assays. According to a white paper published by the vendor, this new product is compatible with Olink Explore 3072(29). Consequently, we believe the research conducted in this study remains relevant and applicable to the Explore HT platform.

## Conclusion

In this study we validate the use of CSF with Olink Explore 3072, however we suggest interpreting results from proteins with NPX values below LOD with extreme caution due to the observed high variance. Additionally, we examined the effects of repeated freeze/thaw cycles on CSF samples when analyzed with PEA using NGS readout and we determined that 3-9 freeze/thaw cycles do not have a significant effect on the detectability of the targets.

## List of abbreviations

ALS: Amyotrophic lateral sclerosis
CNS: Central nervous system
CSF: Cerebrospinal fluid
CV: Coefficient of variation
PEA: Proximity Extension Assay
LOD: Limit of detection
NPX: Normalized protein expression
NGS: Next Generation Sequencing
F/T: freeze/thaw

## Acknowledgments

The authors would like to acknowledge Dong Hyun and Tanmay Shekhar for assisting with the generation of the figures and legends.

## Author’s contributions

MH, MC, XC, AV, HAH conceived of and designed the experiment; MC acquired samples; MH generated data; CL analyzed the data; CL, MH, MC, AV, and HAH interpreted the data; MH, MC, CL prepared the manuscript; MC, MC, CC, AV, HAH, CL reviewed the manuscript.

## Funding

AbbVie funded this study and participated in the study design, research, data collection, analysis, interpretation of data, reviewing, and approval of the abstract. All authors had access to relevant data and participated in the drafting, review, and approval of this abstract. No honoraria or payments were made for authorship.

**Availability of data—The data set analyzed in the current study is included in this article as an additional file**.

## Ethics approval and consent to participate

Informed consent for research use was obtained from all CSF collection subjects by PrecisionMed who owns all necessary approvals and consents.

## AbbVie Disclosure statement

M.H, M.C, A.V, H.A.H, C.L. are employees of AbbVie. X.C is an employee of Argo Biopharma. The design, study conduct, and financial support for this research were provided by AbbVie. AbbVie participated in the interpretation of data, review, and approval of the publication. No honoraria or payments were made for authorship.

## Author details

Genomics Research Center AbbVie, Inc.

1 N Waukegan Rd, AP31-1 North Chicago, IL 60064

Argo Biopharma,

581 Shenkuo Road, 4th Floor

Zhongtong Building, Pudong New Area, Shanghai

