## Supplementary data for "Validation and pre-analytical considerations for processing cerebrospinal fluid samples on a high-throughput proximity extension assay platform"

**Supplementary Table S1. Median Coefficient of Variation per Panel (NPX > LOD)**

| Panel | 1 | 2 | 3 | 4 | 5 | 6 | Mean |
|---|---|---|---|---|---|---|---|
| Cardiometabolic-II | 0.14 | 0.08 | 0.07 | 0.10 | 0.14 | 0.14 | 0.11 |
| Cardiometabolic | 0.08 | 0.13 | 0.11 | 0.09 | 0.16 | 0.12 | 0.12 |
| Inflammation-II | 0.10 | 0.10 | 0.07 | 0.07 | 0.14 | 0.12 | 0.10 |
| Inflammation | 0.10 | 0.12 | 0.10 | 0.12 | 0.19 | 0.13 | 0.12 |
| Neurology-II | 0.13 | 0.10 | 0.08 | 0.11 | 0.12 | 0.15 | 0.11 |
| Neurology | 0.10 | 0.12 | 0.12 | 0.12 | 0.18 | 0.12 | 0.13 |
| Oncology-II | 0.14 | 0.11 | 0.10 | 0.13 | 0.13 | 0.13 | 0.12 |
| Oncology | 0.08 | 0.10 | 0.11 | 0.13 | 0.18 | 0.13 | 0.12 |

**Supplementary Table S2. Proteins Sensitive to Freeze/Thaw cycling from Previously Described Experiments[1]**

| *Protein* | *UniProt ID* | *Gene* | *Explore 384 panel* | *Sensitive to* | *Effect* | *Original Platform* |
|---|---|---|---|---|---|---|
| *Transthyretin* | P02766 | TTR | INFII | Freeze/thaw (1 cycle) | Decreased | SELDI-TOF MS |
| *Cathepsin D* | P07339 | CTSD | CAR | Freeze/thaw (5 cycles) | Decreased | Fluorimetric assays |
| *Protein kinase B alpha/beta/gamma (RAC family)* | Q9Y243 | AKT3 | ONC | Freeze/thaw (8 cycles) | Decreased | Olink |
| *C-type lectin domain family 11 member A* | Q9Y240 | CLEC11A | NEU | Freeze/thaw (8 cycles) | Decreased | Olink |
| *Lipopolysaccharide-binding protein* | P18428 | LBP | CAR | Freeze/thaw (8 cycles) | Decreased | SOMAscan |
| *α-Fucosidase* | P04066 | FUCA1 | CAR | Freeze/thaw (2 cycles) | Increased | Fluorimetric assays |
| *α-Mannosidase* | Q9BV94 | EDEM2 | CARII | Freeze/thaw (1 cycle) | Decreased | Fluorimetric assays |
| | O60476 | MAN1A2 | CARII | | | |
| | Q9Y2E5 | MAN2B2 | CARII | | | |
| | Q5VSG8 | MANEAL | ONCII | | | |
| *β-Galactosidase* | P16278 | GLB1 | NEU | Freeze/thaw (5 cycles) | Decreased | Fluorimetric assays |
| | P06280 | GLA | INFII | | | |

[1]*Hok-A-Hin YS, Willemse EAJ, Teunissen CE, Del Campo M. Guidelines for CSF processing and biobanking: impact on the identification and development of optimal CSF protein biomarkers. In: Methods in Molecular Biology. Humana Press Inc.; 2019. p. 27–50.*

**Supplementary Figure S1.**

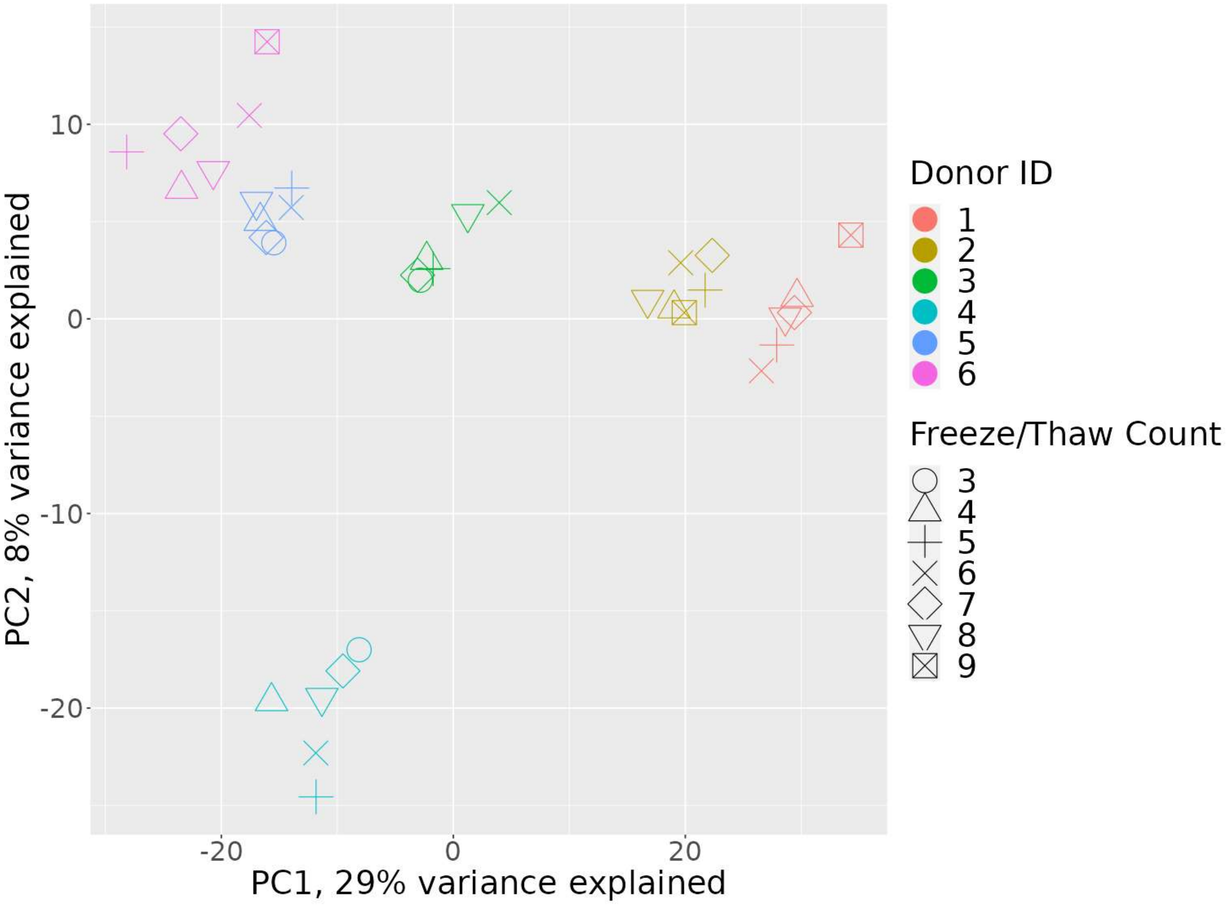


**Supplementary Figure S1. Plot shows principal component 1 (PC1) on x-axis and principal component 2 (PC2) on the y-axis. Color denotes individual donors, and shape denotes the freeze/thaw cycle count. The plot suggests that data points cluster by individual donors, not the freeze/thaw cycle count.**

**Supplementary Figure S2.**

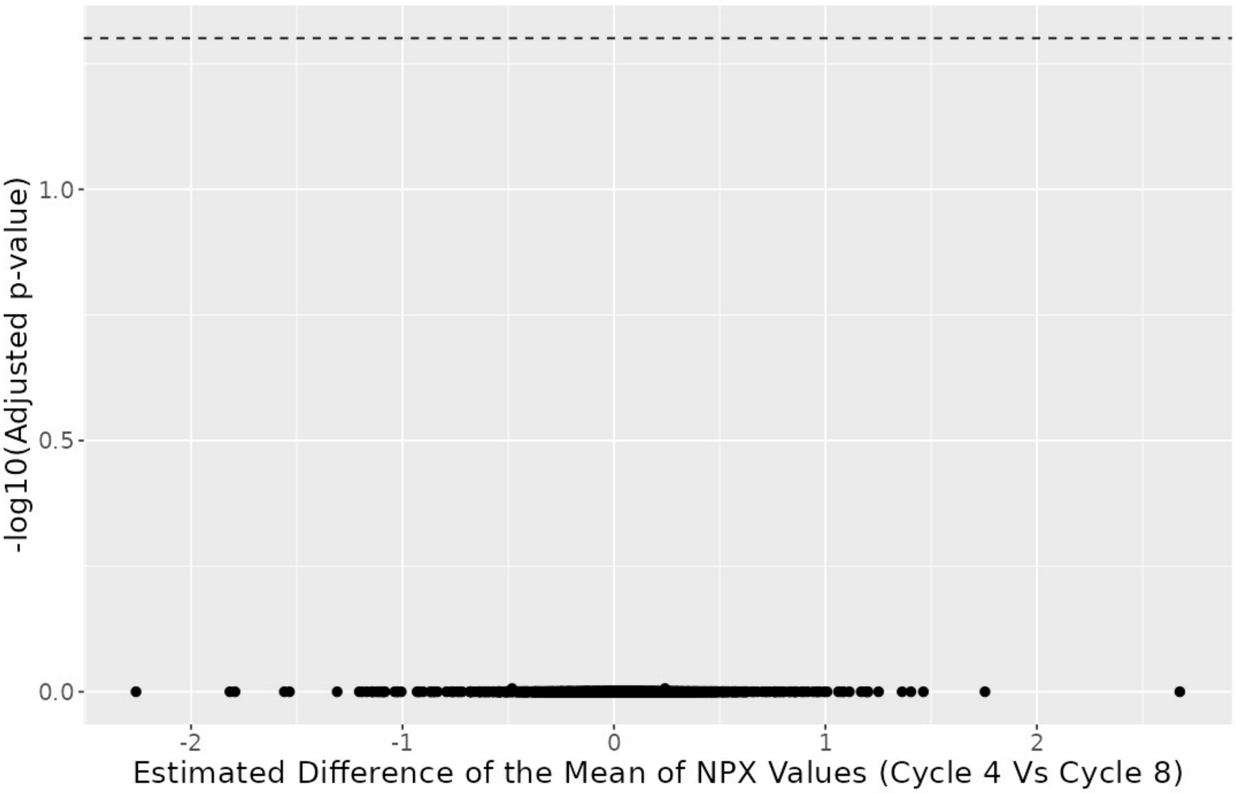


**Supplementary Figure S2. Volcano plot comparing assay levels between freeze/thaw cycle 4 and freeze/thaw cycle 8. Estimated difference of the means of NPX values plotted on the x-axis and adjusted p-value on the y-axis. No significant difference was found for any assay in the study between those cycles. (Min adjusted p value = 0.99)**
